# Nirmatrelvir treatment blunts the development of antiviral adaptive immune responses in SARS-CoV-2 infected mice

**DOI:** 10.1101/2022.11.22.517465

**Authors:** Valeria Fumagalli, Pietro Di Lucia, Micol Ravà, Davide Marotta, Elisa Bono, Stefano Grassi, Lorena Donnici, Rolando Cannalire, Irina Stefanelli, Anastasia Ferraro, Francesca Esposito, Elena Pariani, Donato Inverso, Camilla Montesano, Serena Delbue, Enzo Tramontano, Raffaele De Francesco, Vincenzo Summa, Luca G. Guidotti, Matteo Iannacone

## Abstract

Alongside vaccines, antiviral drugs are becoming an integral part of our response to the SARS-CoV-2 pandemic. Nirmatrelvir – an orally available inhibitor of the 3-chymotrypsin-like cysteine protease – has been shown to reduce the risk of progression to severe COVID-19. However, the impact of nirmatrelvir treatment on the development of SARS-CoV-2-specific adaptive immune responses is unknown. Here, by using a mouse model of SARS-CoV-2 infection, we show that nirmatrelvir administration early after infection blunts the development of SARS-CoV-2-specific antibody and T cell responses. Accordingly, upon secondary challenge, nirmatrelvir-treated mice recruited significantly fewer memory T and B cells to the infected lungs and to mediastinal lymph nodes, respectively. Together, the data highlight a potential negative impact of nirmatrelvir treatment with important implications for clinical management and might help explain the virological and/or symptomatic relapse after treatment completion reported in some individuals.

## Main text

The COVID-19 outbreak, caused by SARS-CoV-2, has resulted in more than 620 million confirmed infections causing greater than 6.5 million deaths worldwide as of October 2022 (https://coronavirus.jhu.edu/map.html). Despite effective COVID-19 vaccines have been developed at an unprecedented pace, a vast number of people are either unwilling or unable to get vaccinated. SARS-CoV-2 is a beta-coronavirus whose RNA genome encodes for two polyproteins, pp1a and pp1ab, and four structural proteins (*1*). The polyproteins are cleaved by the viral papain-like protease (PL^pro^) and by the viral main protease (M^pro^) – the latter also referred to as 3-chymotrypsin-like cysteine protease (3CL^pro^) – to yield nonstructural proteins that are necessary to viral replication (*2, 3*).

Paxlovid, a combination of an orally available M^pro^ inhibitor termed nirmatrelvir (*4*) and ritonavir, has received an Emergency Use Authorization (EUA) from the Food and Drug Administration (FDA) for the treatment of COVID-19 on December 22, 2021. Despite its effectiveness at reducing viral titers and the risk of progressing to severe COVID-19 (*5, 6*), the impact of nirmatrelvir treatment on the development of adaptive immunity to SARS-CoV-2 is unknown. It is also unclear why some patients experience a surprising rebound of viral load and a rapid relapse of COVID-19 symptoms shortly after the completion of an early and effective nirmatrelvir treatment (*7–15*). Viral sequencing indicates that the relapse is not associated with the selection of treatment-resistant mutations, nor due to infection with different SARS-CoV-2 variants (*7, 8*). Whether treatment rebound is due to an impairment in the development of adaptive immunity necessary to complete SARS-CoV-2 clearance remains to be determined.

Here, we set out to assess the impact of nirmatrelvir treatment on the development of antiviral adaptive immunity in a well-characterized mouse model of COVID-19, based on the controlled administration of aerosolized SARS-CoV-2 to K18-hACE2 transgenic mice (*16*).

Nirmatrelvir was synthesized prior to its publication (*4*) via a multistep convergent approach that differed from the reported procedure (*3, 17, 18*). Details and synthesis intermediates are shown in **Figure 1A** and in **Materials and Methods**. As expected, nirmatrelvir was able to inhibit the activity of the SARS-CoV-2 M^pro^ in a dose-dependent manner, as determined by a fluorescence resonance energy transfer (FRET)-based biochemical assay performed both by preincubating the protease with the compound for 30 minutes at 37°C (**Figure 1B**) or by directly adding the substrate to the reaction (**Figure S1A**), as described (*4*). In the two assays, the 50% inhibitory concentration (IC_50_) of nirmatrelvir on the M^pro^ was 47 nM (**Figure 1B**) and 14 nM (**Figure S1A**), respectively. In both procedures, we used the commercially available SARS-CoV-2 M^pro^ inhibitor GC376 (*17–20*) as positive control, showing an IC_50_ of 0.14 nM (**Figure 1B**) and 4.8 nM (**Figure S1A**), respectively. Next, the antiviral activity of nirmatrelvir against SARS-CoV-2 was examined by monitoring the protection from cytopathic effect in infected HEK293T-hACE2 cells (**Figure 1C**) and by quantifying SARS-CoV-2 RNA in the supernatant (**Figure S1B**). Nirmatrelvir prevented death of HEK293T-hACE2 cells infected with the SARS-CoV-2 variants D614G, B.1.617.2 (Delta) and B.1.1.529 (Omicron BA.1) with a mean IC_50_ value of 33 ± 10 nM (**Figure 1C**) and prevented SARS-CoV-2 RNA release with a mean IC_50_ value of 54 ± 25 nM (**Figure S1B**). Finally, we evaluated the antiviral activity of nirmatrelvir in a well-characterized mouse model of COVID-19, based on the controlled administration of aerosolized SARS-CoV-2 to K18-hACE2 transgenic mice (*16*). Briefly, non-anesthetized mice were placed in a nose-only inhalation tower system and exposed to a target dose of 2 × 10^5^ tissue culture infectious dose 50 (TCID_50_) of aerosolized B.1.1.529 under controlled pressure, temperature, and humidity conditions (**Figure 1D**). Mice were treated for six times with vehicle or nirmatrelvir via oral gavage (150 mpk/mouse) starting at 4 hours post infection (p.i.) and every 12 hours thereafter up until day 3 p.i. (**Figure 1D**). The plasma concentration of nirmatrelvir evaluated 4 hours after the last administration was 1.39 ± 0.73 μM (**Figure S1C**). As expected, neither SARS-CoV-2 infection nor nirmatrelvir treatment affected the body weight of K18-hACE2 mice (**Figure 1E**). In line with previously published studies (*4, 16*), whereas vehicle-treated SARS-CoV-2 infected mice showed robust viral replication in the lungs and in the nasal turbinates, nirmatrelvir-treated SARS-CoV-2 infected mice showed virtually undetectable viral RNA and infectious virus in the same anatomical compartments (**Figure 1F, G**).

**Figure 1.**
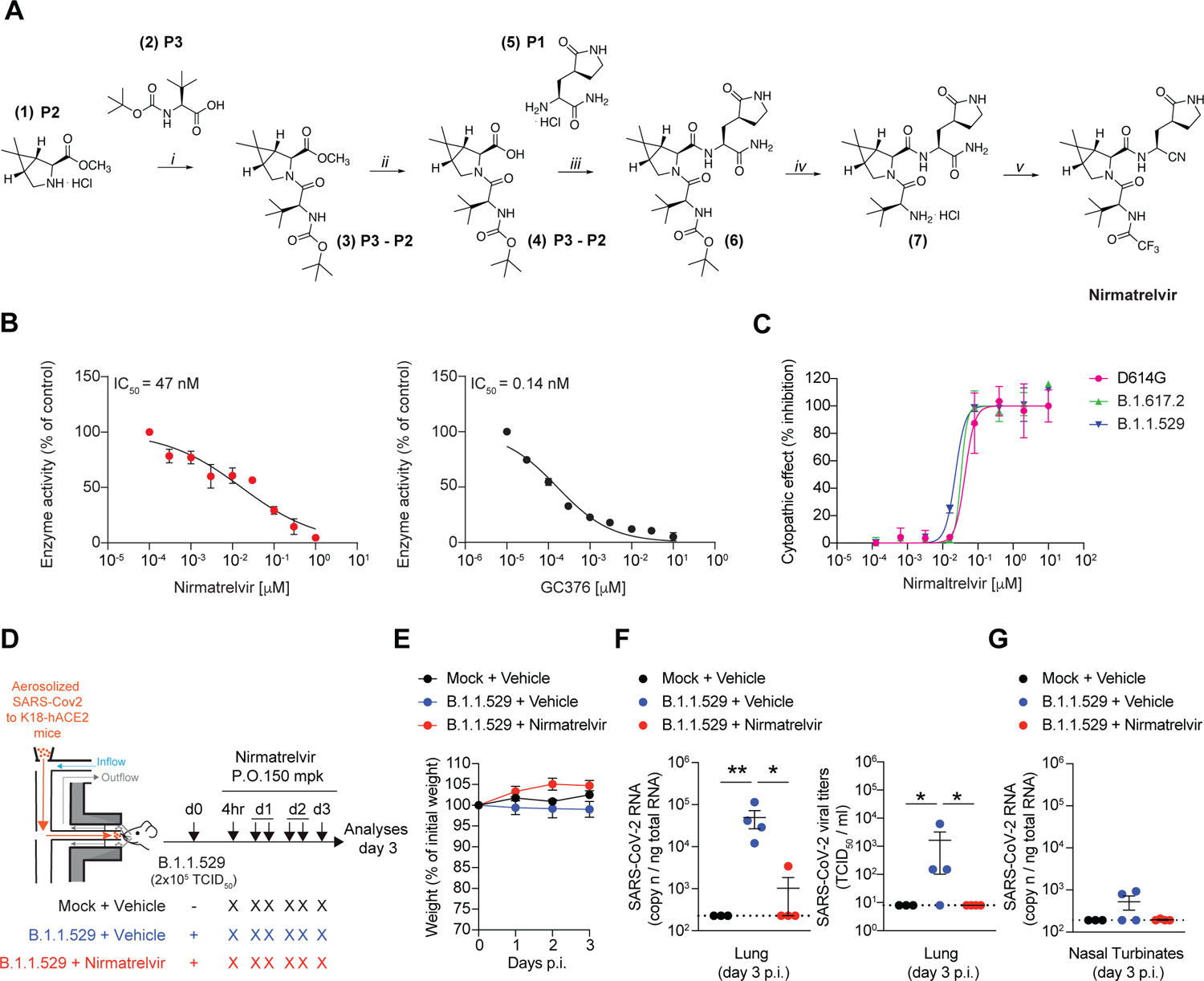
Synthesis of nirmatrelvir and characterization of its biochemical and antiviral activity. **(A)** Synthesis of nirmatrelvir. Reagents and conditions: *i*) HBTU, DIPEA, dry CH_2_Cl_2_, room temperature (RT), 16h, 78%; *ii*) 1N aq. LiOH/THF (1:1), RT, 2h, 100%; *iii)* HBTU, DIPEA, dry CH_2_Cl_2_/DMF, RT, 3h, 75%; *iv)* 4N HCl in 1,4-dioxane/CH_2_Cl_2_ (1:1), 0°C to RT, 2h, 100%; *v*) a) TFAA, dry Py, dry CH_2_Cl_2_, 0°C to RT, 2h; b) TFAA, dry Py, dry CH_2_Cl_2_, 0°C to RT, 15h, 40% over a and b. **(B)** Dose-dependent inhibition of nirmatrelvir (left panel) and GC376 (right panel) on SARS-CoV-2 M^pro^. Prior to adding the substrate for the biochemical reaction, the protease was preincubated for 30 minutes at 37°C with the indicated concentrations of the compound. Data are represented as mean ± SD. **(C)** Dose-dependent antiviral activity of nirmatrelvir in HEK293T-hACE2 cells infected with SARS-CoV-2 D614G (purple symbols), B.1.617.2 (green symbols) or B.1.1.529 (blue symbols). Antiviral activity was determined as percent inhibition of the virus-induced cytopathic effect. Data are represented as mean ± SD. **(D)** Schematic representation of the experimental set up. Non-anesthetized K18-hACE2 transgenic mice were infected with a target dose of 2 x 10^5^ TCID_50_ of SARS-CoV-2 B.1.1.529 through aerosol exposure (see **Materials and Methods** for details). Infected mice were treated with 150 mg/kg (mpk) of nirmatrelvir (red symbols, *n* = 4) or vehicle (blue symbols, *n* = 4) for six times by oral gavage (P.O.) starting 4 hours post infection (p.i.), and every 12 hours thereafter. Mock-treated mice were used as control (black symbols, *n* = 3). Lung, nasal turbinates, and blood were collected and analyzed 3 days p.i. **(E)** Mouse body weight was monitored daily and is expressed as the percentage of weight relative to the initial weight. **(F)** Quantification of SARS-CoV-2 RNA (left panel) and viral titers (right panel) in the lung 3 days after infection. RNA values are expressed as copy number per ng of total RNA and the limit of detection is indicated as a dotted line. Viral titers were determined by median tissue culture infectious dose (TCID_50_). **(G)** Quantification of SARS-CoV-2 RNA in the nasal turbinates 3 days after infection. RNA values are expressed as copy number per ng of total RNA and the limit of detection is indicated as a dotted line. Data in **E**-**G** are represented as mean ± SEM. Data are representative of at least 2 independent experiments. * p-value < 0.05, ** p-value < 0.01; two-way ANOVA followed by Sidak’s multiple comparison test (**E**); Kruskal-Wallis test followed by uncorrected Dunn’s test, each comparison stands alone (**F**, **G**).

With this established system, we next set out to study the consequences of nirmatrelvir treatment on antiviral immune responses. To this end, we infected and treated another group of K18-hACE2 transgenic mice exactly as before but monitored them until 24 days p.i. to assess the SARS-CoV-2 specific antibody response in the sera. Because SARS-CoV-2 T cells are not readily detectable in the blood of K18-hACE2 transgenic mice infected with B.1.1.529 (**Figure S2A, B**), we decided to subject the mice to a homologous re-challenge with a higher dose (1 × 10^6^ TCID_50_) of aerosolized SARS-CoV-2 B.1.1.529 to evaluate the eventual recruitment of memory T (and B) cells to the infected lung and lung-draining mediastinal lymph nodes. The mean plasma concentration of nirmatrelvir 4 hours after the last administration was 1.40 ± 0.99 μM (**Figure S2C**), remarkably similar to the previous experiment. No mice exhibited significant weight loss for the whole duration of the experiment (**Figure 2B**). Of note, the levels of total IgG specific for the spike S1 subunit (Receptor Binding Domain, RBD) (**Figure 2C**) and the levels of anti-B.1.1.529 neutralizing antibodies (**Figure 2D**) were remarkably reduced in nirmatrelvir-treated mice 14 and 21 days p.i., respectively, and 4 days after re-challenge (**Figure 2E, F**). Consistent with this, B cells recovered from the mediastinal lymph-nodes of nirmatrelvir-treated mice 4 days after re-challenge exhibited a lower expression of the activation marker CD95 (**Figure 2G**) and in one vehicle-treated mouse we could detect RBD-specific B cells (**Figure 2H**). Additionally, large lymphocytic aggregates consisting of proliferating B cells were detected in the lungs of vehicle- but not nirmatrelvir-treated mice, 4 days after re-challenge (**Figure S3**). SARS-CoV-2-specific CD8^+^ and CD4^+^ T cells recovered from lung homogenates were assessed for intracellular IFN-γ and TNF-α expression upon *in vitro* stimulation with a pool of SARS-CoV-2 peptides covering the complete nucleocapsid, matrix, and spike proteins (*21*). In line with the results obtained for the humoral response, we found that the frequency and absolute number of IFN-γ^+^ and IFN-γ^+^ TNF-α^+^ SARS-CoV-2-specific CD8^+^ and CD4^+^ T cells were significantly lower in the lungs of nirmatrelvir-treated mice compared to vehicle-treated mice, 4 days after homologous re-challenge (**Figure 2I-N**). Humoral and cellular responses to SARS-CoV-2 were also reduced when nirmatrelvir treatment was initiated later (24 or 48 hours after infection) and when infection was performed with different SARS-CoV-2 variants (**Figure S4**). However, nirmatrelvir treatment of mice infected with unrelated viruses (i.e., vesicular stomatitis virus [VSV] and lymphocytic choriomeningitis virus [LCMV]) did not inhibit the development of antiviral adaptive immune responses, indicating that nirmatrelvir is not *per se* an immune suppressive drug (**Figure S5**).

**Figure 2.**
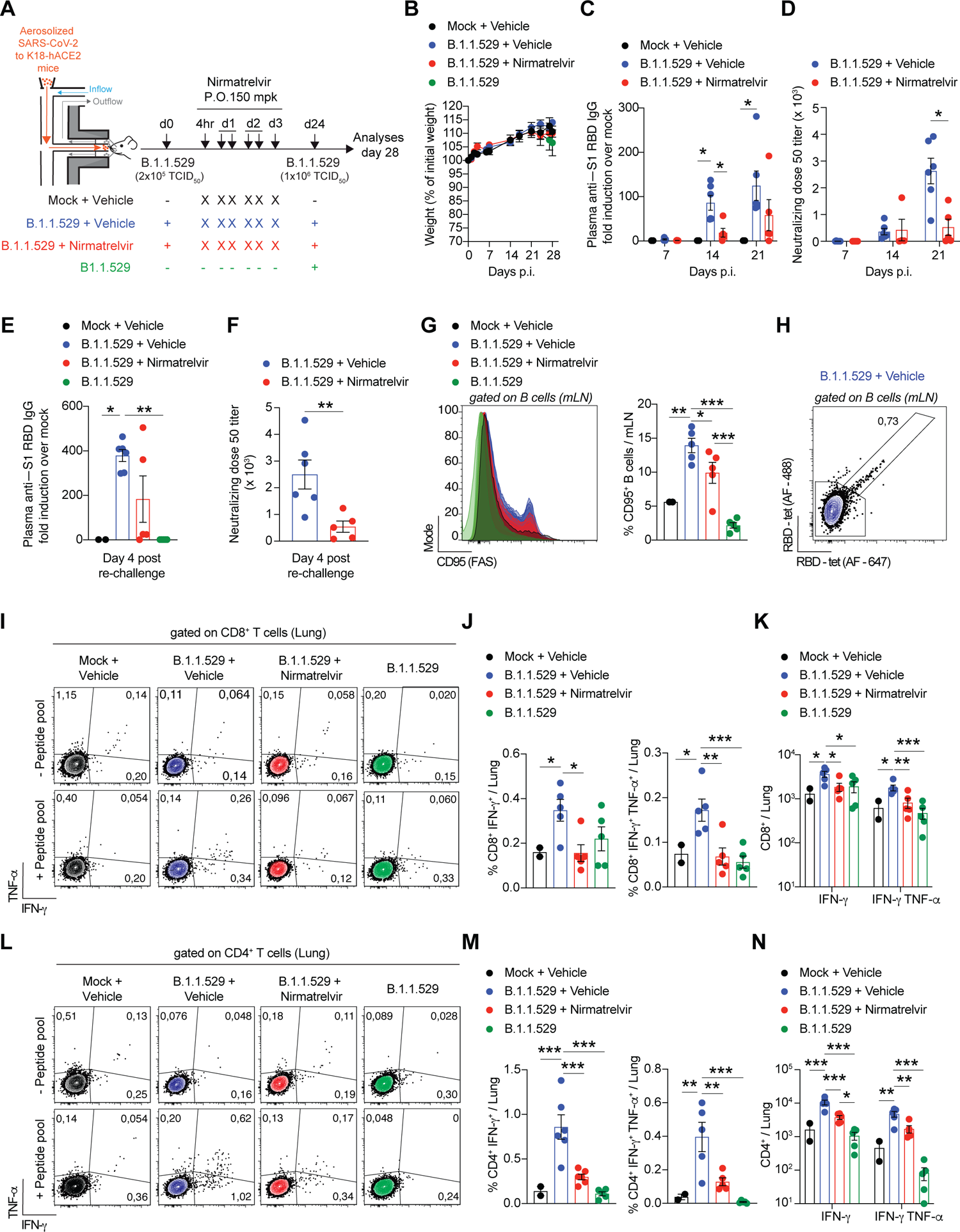
Nirmatrelvir treatment of SARS-CoV-2 infected mice blunts the development of antiviral adaptive immune responses. **(A)** Schematic representation of the experimental set up. Non-anesthetized K18-hACE2 mice were exposed to a target dose of 2 x 10^5^ TCID_50_ of aerosolized SARS-CoV-2 B.1.1.529 (see **Materials and Methods** for details). Infected mice were treated with 150 mpk of nirmatrelvir (red symbols, *n* = 5) or vehicle (blue symbols, *n* = 6) for six times by oral gavage (P.O.) starting 4 hours p.i., and every 12 hours thereafter. Twenty-four days after infection, mice were re-challenged with a target dose of 1 x 10^6^ TCID_50_ of SARS-CoV-2 B.1.1.529 through aerosol exposure. Mock-treated mice were used as non-infected controls (black symbols, *n* = 3). A group of naïve mice challenged with 1 x 10^6^ TCID_50_ of SARS-CoV-2 B.1.1.529 served as additional controls (green symbols, *n* = 5). Blood was collected 7, 14, 21 after the first infection and 4 days post re-challenge. Lung, nasal turbinates, and lung-draining mediastinal lymph nodes (mLN) were collected and analyzed 4 days post re-challenge. **(B)** Mouse body weight was monitored daily and is expressed as percentage of weight relative to the initial weight. **(C, E)** Quantification of anti-S1 RBD IgG levels by ELISA in the plasma of the indicated mice (**C**) 7, 14 and 21 days p.i. or (**E**) 4 days post re-challenge. **(D, F)** Neutralization dose 50 (ND50) against SARS-CoV-2 B.1.1.529 pseudovirus in the plasma of the indicated mice (**D**) 7, 14 and 21 days p.i. or (**F**) 4 days post re-challenge. **(G)** Flow cytometry histogram (left panel) and percentage (right panel) of B cells that stained positive for CD95 in the mLN of the indicated mice 4 days post re-challenge (pre-gated on live^+^/CD4^-^/ CD8^-^/ B220^+^/CD19^+^ cells). **(H)** Flow cytometry plot of RBD-specific B cells detected by two fluorescently labeled streptavidin-based RBD-biotinylated tetramers in the mLN of one vehicle-treated mouse 4 days post re-challenge (pre-gated on live^+^/CD4^-^/ CD8^-^/ B220^+^/CD19^+^ cells). **(I, L)** Representative flow cytometry plots of (**I**) CD8^+^ T cells or (**L**) CD4^+^ T cells expressing IFN-γ and TNF-α in the lungs of the indicated mice 4 days post re-challenge. Unstimulated cells are shown in the upper panels, whereas cells re-stimulated with a pool of SARS-CoV-2 peptides for 4 hours at 37°C are shown in the bottom panels. Plots were pre-gated as (**I**) live^+^/ B220^-^/CD19^-^/CD4^-^/CD8^+^ cells or (**L**) live^+^/B220^-^/CD19^-^/CD8^-^/CD4^+^. **(J, K, M, N).** Frequency (**J, M**) and absolute number (**K, N**) of IFN-γ− and TNF-α−producing CD8^+^ T cells (**J, K**) or CD4^+^ T cells (**M, N**) in the lung of the indicated mice 4 days post re-challenge. Data are expressed as mean ± SEM and are representative of at least 2 independent experiments. * p-value < 0.05, ** p-value < 0.01, *** p-value < 0.001; two-way ANOVA followed by Sidak’s multiple comparison test (**B-D**); Kruskal-Wallis test followed by uncorrect Dunn’s test, each comparison stands alone (**E**); Mann-Whitney test (**F**); One-way ANOVA followed by uncorrected Fisher’s LSD, each comparison stands alone (**G, J, K, M, N**). Normal distribution was verified by Shapiro-Wilk test.

Of note, we did not detect viral titers in the lungs of all but one nirmatrelvir-treated mouse after homologous re-challenge (**Figure S6A, B**). Similarly, infection of K18-hACE2 transgenic mice with the poorly pathogenic Omicron (B.1.1.529) variant (*22*) did not allow us to evaluate clinical signs of disease. We believe that the failure to observe decreased protection upon re-infection in mice treated with nirmatrelvir (despite a prominent reduction in adaptive immunity) has to do with the limitations of the experimental setup. Case in point, we infected mice with Omicron (B.1.1.529) variant, we treated or not with nirmatrelvir and we re-challenged mice with Delta (B1.1.617) variant, known to be more pathogenic and to replicate at higher levels in mice. As shown in **Figure S6C, D**, two out of six mice that were treated with nirmatrelvir during the primary infection showed higher viral RNA in the lungs upon Delta re-challenge. Future studies with different SARS-CoV-2 variants and/or doses should assess the impact of the reduced adaptive immunity caused by nirmatrelvir treatment on viral load and the severity of disease progression upon re-challenge.

In summary, our results indicate that nirmatrelvir treatment early after infection negatively impacts the development of adaptive immune response to SARS-CoV-2 in K18-hACE2 transgenic mice. Although the mechanistic bases behind this observation were not addressed in this study, it is conceivable that this is due to insufficient antigen exposure (quantity and/or duration) of naïve B and T cells. It is worth noting that successful antimicrobial treatment does not inevitably result in reduced adaptive immune responses to any pathogen. For instance, treatment of mice infected with *Listeria monocytogenes* with amoxicillin early after infection did not significantly impair the development of T cell responses (*23, 24*). Furthermore, treatment with antibiotics before *L. monocytogenes* infection allowed the development of functional antigen-specific memory CD8^+^ T cells in the absence of contraction (*25*). Thus, the effect of antiviral therapy on adaptive immunity probably depends on the impact of such treatment on several factors including not only pathogen replication but also duration of antigen expression and presentation, activation of innate immunity, etc. While the clinical data continue to support nirmatrelvir treatment for the prevention of severe COVID-19 in high-risk individuals (*5*), the data reported here draw attention to a potential negative impact of this therapy. Whether this effect is an exclusive feature of nirmatrelvir or whether forthcoming antivirals acting on SARS-CoV-2 would have similar effects should be addressed in future studies. Although mice do not reproduce the viral rebound observed in some patients treated with nirmatrelvir, we believe that our results might help explain the virological and/or symptomatic relapse after treatment completion reported in some individuals and should inform clinical and public health policies.

## Acknowledgments

We thank S. De Palma, A. Fiocchi, L. Giustini, M. Mainetti, C. Perucchini for technical support; M. Silva for secretarial assistance; L. Aurisicchio (Takis Biotech) for providing biotinylated-RDB, and the members of the Iannacone laboratory for helpful discussions. Flow cytometry was carried out at FRACTAL, a flow cytometry resource and advanced cytometry technical applications laboratory established by the San Raffaele Scientific Institute. Confocal immunofluorescence histology was carried out at Alembic, an advanced microscopy laboratory established by the San Raffaele Scientific Institute and the Vita-Salute San Raffaele University. We would like to acknowledge the PhD program in Basic and Applied Immunology and Oncology at Vita-Salute San Raffaele University, as D.M. conducted this study as partial fulfillment of their PhD in Molecular Medicine within that program. M.I. is supported by the European Research Council (ERC) Consolidator Grant 725038, ERC Proof of Concept Grant 957502, Italian Association for Cancer Research (AIRC) Grants 19891 and 22737, Italian Ministry of Health Grants RF-2018-12365801, and Funded Research Agreements from Gilead Sciences, Asher Bio, Takis Biotech and Vir Biotechnology. L.G.G. is supported by the Italian Association for Cancer Research (AIRC) Grant 22737, Lombardy Open Innovation Grant 229452, PRIN Grant 2017MPCWPY from the Italian Ministry of Education, University and Research, Funded Research Agreements from Gilead Sciences, Avalia Therapeutics and CNCCS SCARL and a donation from FONDAZIONE SAME for COVID-19-related research. V.F. is supported by a donation from FONDAZIONE PROSSIMO MIO and from AIRC Fellowship for Italy (ID 26813-2021).

## Author contributions

Conceptualization, V.F., L.G.G., M.I.; Investigation, V.F., P.D.L., M.R., D.M., E.B., C.P., S.G., L.D., R.C., I.S., A.F., F.E., E.P., D.I., C.M.; Resources, L.D., R.C., I.S., A.F., S.D., R.D.F., V.S.; Formal Analysis, V.F., L.D., R.C., I.S., A.F., F.E., E.P., E.T., R.D.F., V.S.; Writing V.F., M.I. with input from all authors; Visualization, V.F.; Funding Acquisition, L.G.G. and M.I.; Project supervision, M.I.

## Data and materials availability

All data are available in the main text or in the supplementary material.

## Supplementary Figure Legends

**Figure S1.**
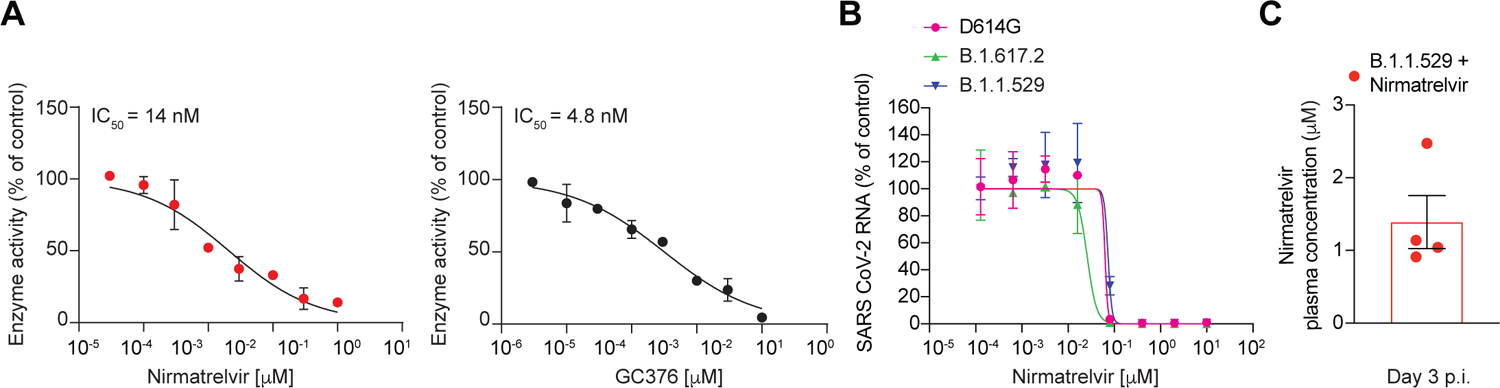
Biochemical activity, antiviral efficacy, and plasma concentrations of nirmatrelvir. **(A)** Dose-dependent inhibition of nirmatrelvir (left panel) and GC376 (right panel) on SARS-CoV-2 M^pro^. The enzymatic reaction was immediately initiated with the addition of the substrate. Data are represented as mean ± SD. **(B)** Dose-dependent antiviral activity of nirmatrelvir in HEK293T-hACE2 cells infected with SARS-CoV-2 D614G (purple symbols), B.1.617.2 (green symbols) and B.1.1.529 (blue symbols). Antiviral activity was determined by qPCR quantification of SARS-CoV-2 RNA in the supernatant. Data are represented as mean ± SD. **(C)** Nirmatrelvir concentration (μM) in the plasma of mice treated as described in Figure 1D. Data are represented as mean ± SEM and are representative of at least two independent experiments.

**Figure S2.**
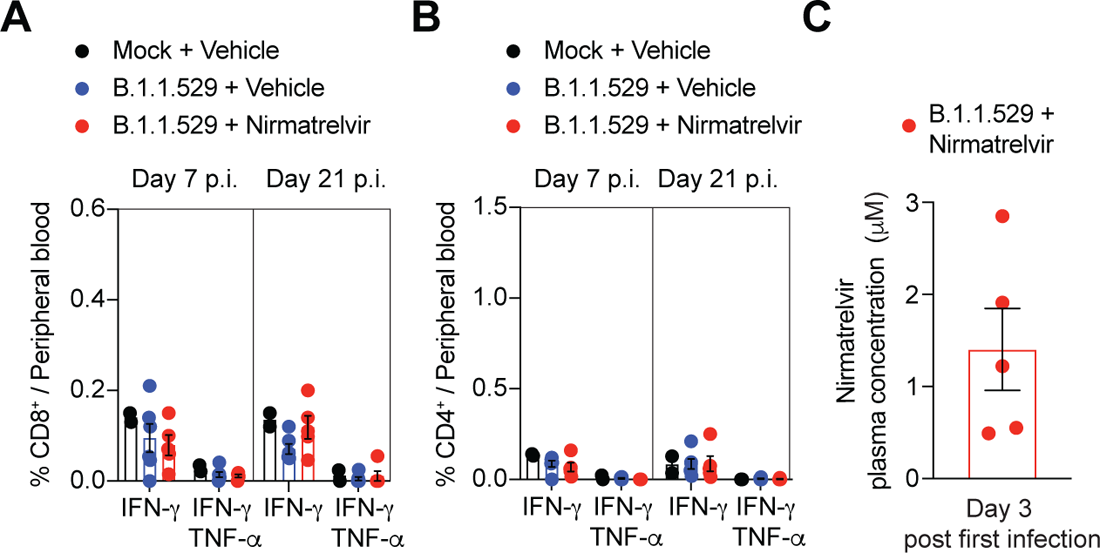
Absence of detectable SARS-CoV-2 specific T cells in the blood of K18-hACE2 transgenic mice infected with B.1.1.529. **(A, B)** Frequency of IFN-γ− and TNF-α−producing CD8^+^ T cells (**A**) or CD4^+^ T cells (**B**) in the peripheral blood of the indicated mice 7 and 21 days after the first infection. Cells were stimulated *in vitro* with a pool of SARS-CoV-2 peptides for 4 hours at 37°C. Plots were pre-gated as (**A**) live^+^/ B220^-^/CD19^-^/CD4^-^/CD8^+^ cells or (**B**) live^+^/B220^-^/CD19^-^/CD8^-^/CD4^+^. **(C)** Nirmatrelvir concentration (μM) in the plasma of mice treated as described in Figure 2A, 3 days after the first infection. Data are represented as mean ± SEM and are representative of at least two independent experiments.

**Figure S3.**
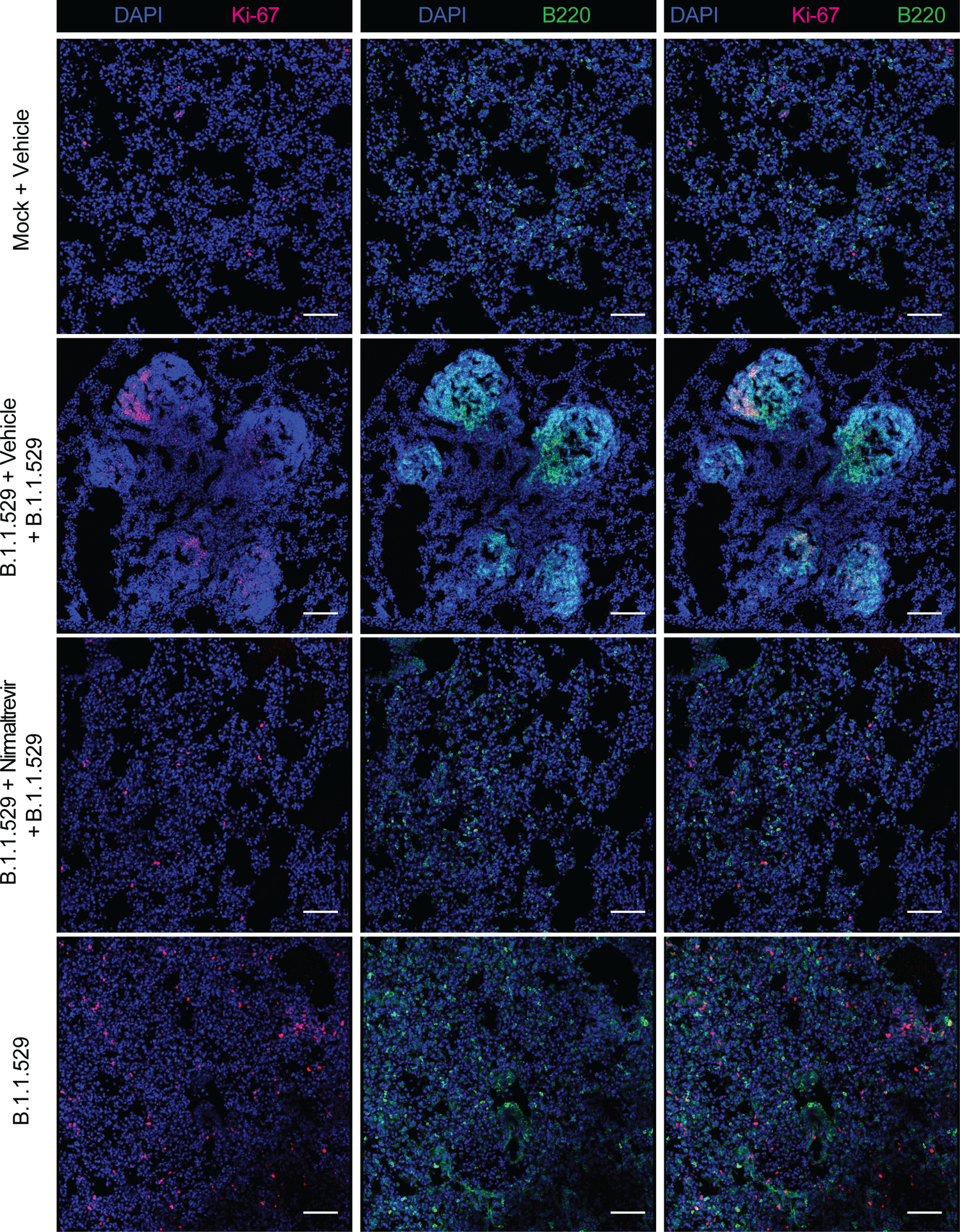
Lymphocytic aggregates in the lungs of vehicle- but not nirmatrelvir-treated mice. Representative confocal immunofluorescence micrographs of lung sections from mice 4 days post re-challenge, as described in Figure 2A. Mock-treated mice (first lane), vehicle-treated mice (second lane), nirmatrelvir-treated mice (third lane) and naïve-challenged mice (fourth lane). Ki-67^+^ cells are depicted in purple; B220^+^ cells are depicted in green and cell nuclei in blue. Scale bars, 50 μm.

**Figure S4.**
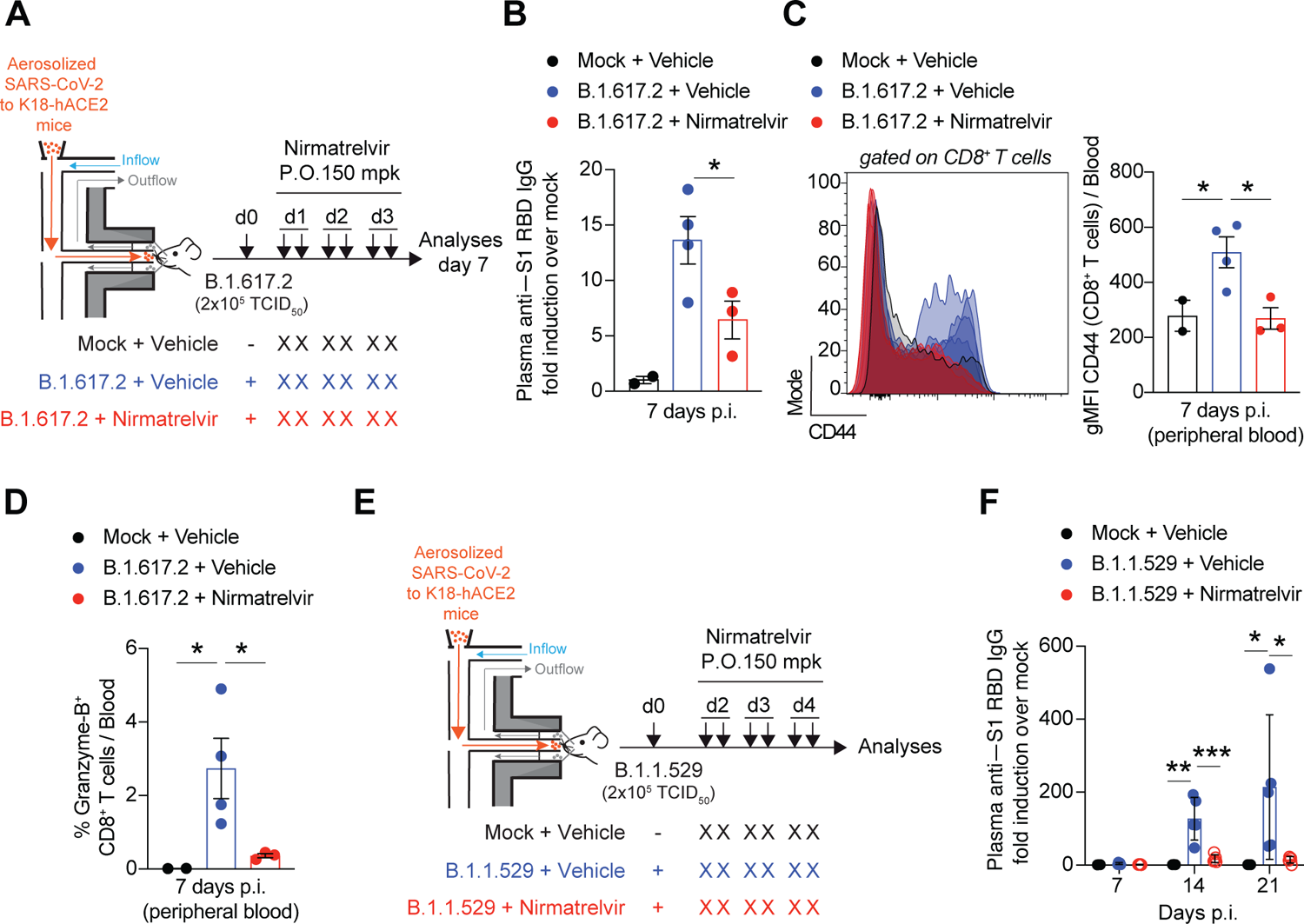
Adaptive immune responses are blunted in mice treated with nirmatrelvir 24 or 48 hours after SARS-CoV-2 infection. **(A)** Schematic representation of the experimental set up. Non-anesthetized K18-hACE2 mice were exposed to a target dose of 2 x 10^5^ TCID_50_ of aerosolized SARS-CoV-2 B.1.617.2. Infected mice were treated with 150 mpk of nirmatrelvir (red symbols, *n* = 3) or vehicle (blue symbols, *n* = 4) for six times by oral gavage (P.O.) starting 24 hours p.i., and every 12 hours thereafter. Mock-treated mice were used as non-infected controls (black symbols, *n* = 2). **(B)** Quantification of anti-S1 RBD IgG levels by ELISA in the plasma of the indicated mice 7 days p.i.. **(C)** Flow cytometry histogram (left panel) and geometric mean fluorescent intensity (gMFI) quantification (right panel) of CD44 expression by CD8^+^ T cells in the blood of the indicated mice 7 days p.i. (pre-gated on live^+^/ B220^-^/ CD19^-^/ CD4^-^/ CD8^+^ cells). **(D)** Frequency of Granzyme-B−producing CD8^+^ T cells in the blood of the indicated mice 7 days p.i.. Cells were stimulated *in vitro* with a pool of SARS-CoV-2 peptides for 4 hours at 37°C. Plots were pre-gated as (**C**) live^+^/ B220^-^/CD19^-^/CD4^-^/CD8^+^ cells. **(E)** Schematic representation of the experimental set up. Non-anesthetized K18-hACE2 mice were exposed to a target dose of 2 x 10^5^ TCID_50_ of aerosolized SARS-CoV-2 B.1.1.529. Infected mice were treated with 150 mpk of nirmatrelvir (red symbols, *n* = 5) or vehicle (blue symbols, *n* = 5) for six times by oral gavage (P.O.) starting 48 hours p.i., and every 12 hours thereafter. Mock-treated mice were used as non-infected controls (black symbols, *n* = 3). **(F)** Quantification of anti-S1 RBD IgG levels by ELISA in the plasma of the indicated mice 7, 14 and 21 days p.i.. Data are expressed as mean ± SEM. * p-value < 0.05; One-way ANOVA followed by uncorrected Fisher’s LSD, each comparison stands alone. Normal distribution was verified by Shapiro-Wilk test.

**Figure S5.**
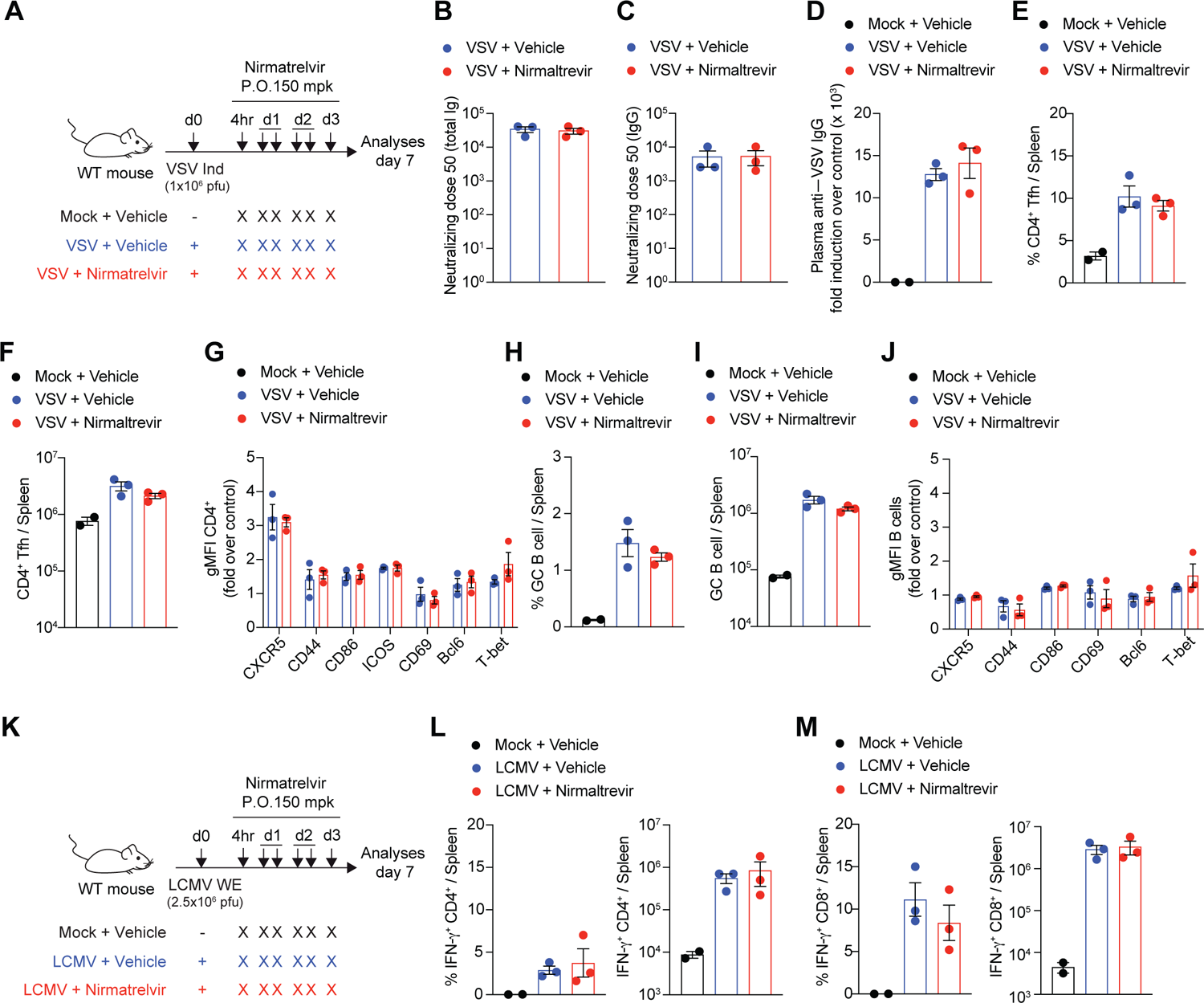
Nirmatrelvir treatment does not impair the development of adaptive immune responses to VSV or LCMV. **(A, K)** Schematic representation of the experimental set up. WT mice were intravenously infected with 1 x 10^6^ pfu of VSV Indiana (**A**) or 2.5 x 10^6^ pfu of LCMV WE (**K**). Infected mice were treated with 150 mpk of nirmatrelvir (red symbols, *n* = 3) or vehicle (blue symbols, *n* = 3) for six times by oral gavage (P.O.) starting 4 hours p.i., and every 12 hours thereafter. Mock-treated mice were used as non-infected controls (black symbols, *n* = 2). Blood and spleen were collected 7 days p.i.. **(B, C)** Neutralization dose 50 (ND50) of total immunoglobulin (Ig) (**B**) or IgG (**C**) against VSV in the plasma of the indicated mice. **(D)** Quantification of anti-VSV IgG levels by ELISA in the plasma of the indicated mice. **(E, F)** Frequency (**E**) and absolute number (**F**) of CD4^+^ T follicular helper cells (Tfh) in the spleen of indicated mice. Tfh cells were defined as live^+^/ B220^-^/ CD19^-^/ CD8^-^/ CD4^+^/ CXCR5^+^/ Bcl6^+^ cells. **(D)** Geometric mean fluorescent intensity (gMFI) of markers expressed by CD4^+^ T cells. Values are showed as fold expression over control. **(H, I)** Frequency (**H**) and absolute number (**I**) of germinal center (GC) B cells in the spleen of indicated mice. GC B cells were defined as live^+^/ CD8^-^/ CD4^-^/ B220^+^/ CD19^+^/ GL7^+^ / Bcl6^+^ cells. **(J)** Geometric mean fluorescent intensity (gMFI) of markers expressed by B cells. Values are showed as fold expression over control. **(L, M)** Frequency (left panel) and absolute number (right panel) of IFN-γ−producing CD4^+^ T cells **(L)** and CD8^+^ T cells **(M)** in the spleen of indicated mice. Cells were stimulated *in vitro* with the LCMV peptides GP61 and GP33 for 4 hours at 37°C. Data are represented as mean ± SEM.

**Figure S6.**
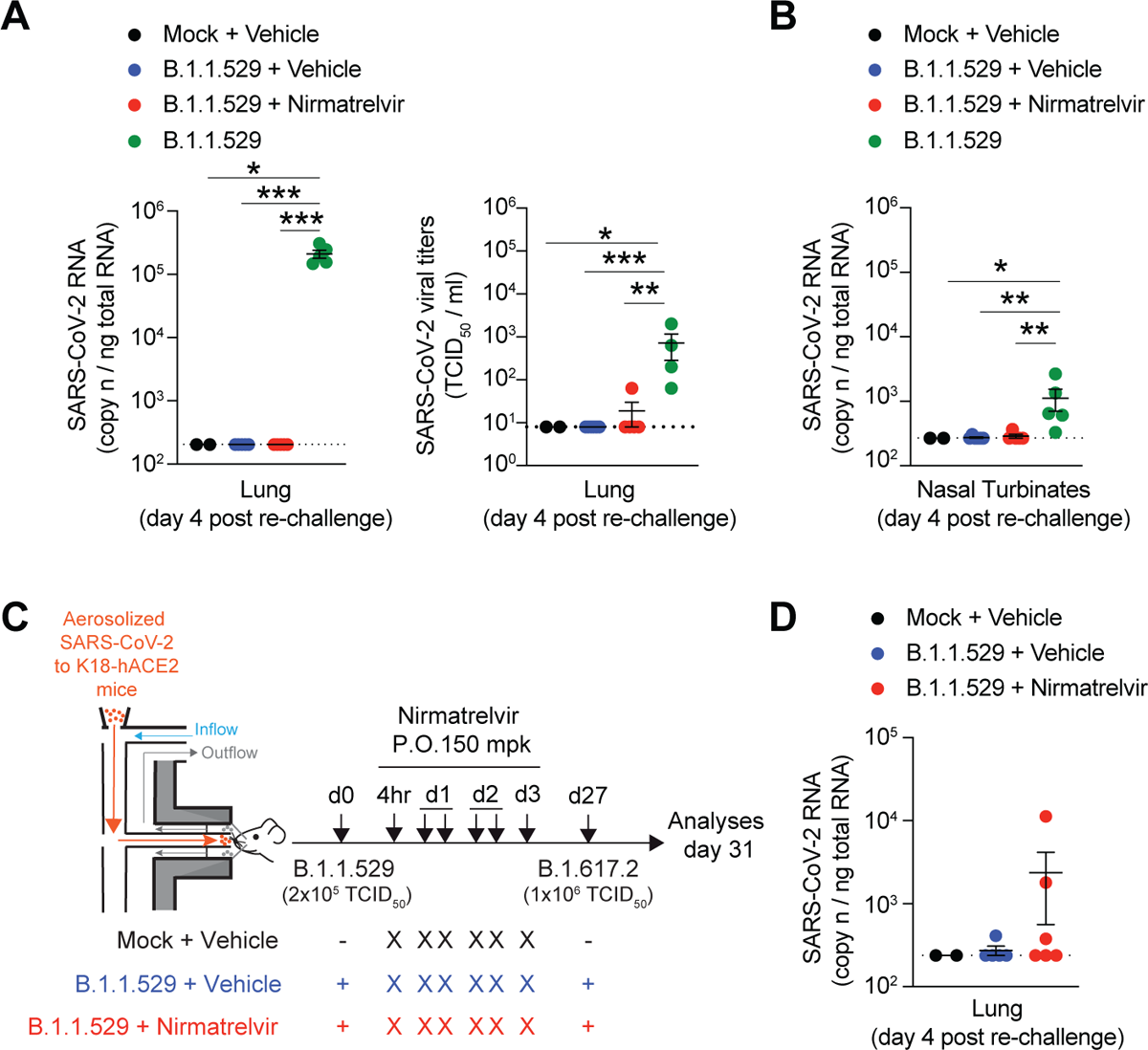
Effect of nirmatrelvir treatment on viral titers upon SARS-CoV-2 re-challenge. **(A)** Quantification of SARS-CoV-2 RNA (left panel) and viral titers (right panel) in the lungs of the indicated mice treated as described in Figure 2A. RNA values are expressed as copy number per ng of total RNA. Viral titers were determined by median tissue culture infectious dose (TCID_50_). Limit of detection is indicated as a dotted line. Blue, vehicle-treated mice; red, nirmatrelvir-treated mice. Mock-treated mice were used as non-infected controls (black symbols). A group of naïve mice challenged with 1 x 10^6^ TCID_50_ of SARS-CoV-2 B.1.1.529 served as additional controls (green symbols). **(B)** Quantification of SARS-CoV-2 RNA in the nasal turbinates of the indicated mice 4 days post re-challenge. RNA values are expressed as copy number per ng of total RNA and the limit of detection is indicated as a dotted line. **(C)** Schematic representation of the experimental set up. Non-anesthetized K18-hACE2 mice were exposed to a target dose of 2 x 10^5^ TCID_50_ of aerosolized SARS-CoV-2 (B.1.1.529). Infected mice were treated with 150 mpk of nirmatrelvir (red symbols, *n* = 6) or vehicle (blue symbols, *n* = 6) for six times by oral gavage (P.O.) starting 4 hours p.i., and every 12 hours thereafter. Twenty-seven days after infection, mice were re-challenged with a target dose of 1 x 10^6^ TCID_50_ of SARS-CoV-2 (B.1.617.2) through aerosol exposure. Mock-treated mice were used as non-infected controls (black symbols, n = 2). **(D)** Quantification of SARS-CoV-2 RNA in the lung 4 days after re-challenge. RNA values are expressed as copy numbers per ng of total RNA and the limit of detection is indicated as a dotted line. Data are represented as mean ± SEM and are representative of at least two independent experiments. * p-value < 0.05, ** p-value < 0.01, *** p-value < 0.001; Kruskal-Wallis test followed by uncorrect Dunn’s test, each comparison stands alone.

## Material and Methods

### Viruses

The SARS-CoV-2 isolates were propagated in Vero E6-hTMPRSS2 cells. Briefly, 3 x 10^6^ Vero E6-hTMPRSS2 cells were plated into T75 flask in DMEM 2% FBS. After 24 hours, cells were inoculated with 0.001 or 0.01 MOI of SARS-CoV-2 D614G (hCoV-19/Italy/LOM-UniMI-vir1/2020; EPI_ISL_58405), SARS-CoV-2 B.1.617.2 (hCoV-19/Italy/LOM-Milan-UNIMI9615/2021, EPI_ISL_3073880) or SARS-CoV-2 B.1.1.529 (hCoV-19/Italy/LOM-19182/2021, EPI_ISL_10898045). Supernatant was collected 48-72 hours later, centrifuged for 5 minutes at 500*g*, aliquoted and stored at −80°C. VSV Indiana and LCMV WE were propagated and quantified as described (*26, 27*).

### Nirmatrelvir synthesis

Reagents and solvents were purchased from commercial sources and used without further purification. Reactions were carried out at room temperature (RT), unless otherwise specified. Moisture-sensitive reactions were performed under a positive pressure of dry nitrogen in oven-dried glassware. Analytical thin-layer chromatography (TLC) on silica gel 60 F254 plates (250 µm thickness) was performed to monitor the reaction progress, using UV and KMNO_4_ as revelation method. Analytical HPLC was performed to monitor the reaction progress and the purity of target compound. Flash chromatography on silica gel (70—230 mesh) and preparative HPLC were performed for purification. All products were characterized by their NMR and MS spectra. (ESI)-MS spectra were performed on a LTQ Orbitrap XL mass spectrometer (Thermo Fisher Scientific) by infusion into the ESI source using MeOH as solvent. ^1^H NMR spectra were recorded in deuterated solvents at 25°C on Bruker Avance NEO 400 MHz and 700 MHz instruments. equipped with a RT-DR-BF/1H-5mm-OZ SmartProbe. Chemical shifts (δ) are reported in part per million (ppm) downfield from tetramethylsilane, using residual solvent signal as the internal reference.

The final compound was characterized by HPLC-MS/MS, using a Dionex ULTIMATE 3000 (Thermo Fisher Scientific) HPLC module and a LTQ XL mass spectrometer with electrospray ionization in positive mode and an Ion-Trap detector. Separation was performed with a Kinetex column C18 Polar column (250 mm × 4.6 mm; particle size 5 μm, Phenomenex, Torrance, CA, USA) at 30°C, using a 17 minutes gradient, 5%[0.1%TFA/CH_3_CN]/95%[0.1%TFA/H_2_O] to 95%[0.1%TFA/CH_3_CN]. Analytical HPLC was performed on Shimatzu-1100 HPLC using a Kinetex C18 column (4.6 mm x 150 mm, 5 μm, 100 Å) with an acetonitrile (0.1% HCOOH) − water (0.1% HCOOH) custom gradient. The purity of the final compound was >95%, as determined by HPLC (UV λ = 200 nm). Preparative HPLC was performed on Shimatzu LC-20AP using a Sunfire C18 column (19 mm x 100 mm, 5 μm, 100 Å) with an acetonitrile (0.1% HCOOH) − water (0.1% HCOOH) custom gradient.

Briefly, two “building blocks”, the acid dipeptide P3-P2 and the residue P1, were first synthesized separately and then assembled to generate the advanced intermediate **6**. **6** was deprotected to obtain intermediate **7** that was converted into the final product Nirmatrelvir with one pot two steps procedure by converting 7 primary amide in P1 to the nitrile, electrophilic “warhead”, and obtaining the trifluoroacetamide, as N terminal capping group.

Methyl (1R,2S,5S)-3-((S)-2-((tert-butoxycarbonyl)amino)-3,3-dimethylbutanoyl)-6,6-dimethyl-3-azabicyclo[3.1.0]hexane-2-carboxylate (**3**). Tert-leucine-OH **2** (1 g, 4.32 mmol) was dissolved in dry CH_2_Cl_2_ (5 mL) and amine hydrochloride **1** (1.15 g, 5.62 mmol), HBTU (1.8 g, 4.75 mmol) and DIPEA (1.5 mL, 8.64 mmol) were added at 0 ° C under a nitrogen atmosphere. The resulting solution was kept under magnetic stirring at RT for 16 hours. Then, the reaction mixture was washed with saturated aq. NaHCO_3_ (x1), 1N HCl (x1), brine (x1), dried over anhydrous Na_2_SO_4_, filtered and concentrated under vacuum. The reaction crude was purified by flash column chromatography (Hexane / EtOAc 8: 2) to obtain a colorless oil (reaction time: 16 hours; yield: 1.38 g, 78%). ^1^H NMR (400 MHz, DMSO-d_6_) δ 6.73 (d, J = 9.3 Hz, 1H), 4.21 (s, 1H), 4.05 (d, J = 9.4 Hz, 1H), 3.93 (d, J = 10.4 Hz, 1H), 3.79 (dd, J = 10.3, 5.3 Hz, 1H), 3.65 (s, 3H), 1.55 - 1.49 (m, 1H), 1.41 (d, J = 7.5 Hz, 1H), 1.35 (s, 9H), 1.01 (s, 3H), 0.93 (s, 9H), 0.85 (s, 3H). MS (ESI) m/z calcd: [M + H]^+^ for C_20_H_35_N_2_O_5_^+^ 383.51, found [M + H]^+^ 383.50.

(1R,2S,5S)-3-((S)-2-((tert-butoxycarbonyl)amino)-3,3-dimethylbutanoyl)-6,6-dimethyl-3-azabicyclo[3.1.0]hexane-2-carboxylic acid (**4**). The methyl ester intermediate **3** (1.35 g 3.6 mmol) was dissolved in THF (18 mL), then 1N aq. LiOH was added (18 mmol, 18 mL), and the reaction mixture was kept under stirring at RT for 3 hours. The reaction mixture was cooled to 0 °C, placed in water / ice, acidified with 1 N HCl to pH = 4, then extracted with EtOAc (x3). Then, the collected organic layers were washed with brine (x1), dried over Na_2_SO_4_, filtered and concentrated under vacuum to afford dipeptide acid **4** as white solid which was used in the following step without further purification (reaction time: 2 hours, yield: 1.30 mg, 100%). ^1^H NMR (400 MHz, DMSO-d_6_) δ 12.54 (s, 1H), 6.67 (d, J = 9.4 Hz, 1H), 4.13 (s, 1H), 4.04 (s, 1H), 3.91 (d, J = 10.4 Hz, 1H), 3.77 (dd, J = 10.2, 5.3 Hz, 1H), 1.54 - 1.46 (m, 1H), 1.40 (s, 1H), 1.35 (s, 9H), 1.01 (s, 3H), 0.93 (s, 9H), 0.84 (s, 3H). MS (ESI) m/z calcd: [M + H]^+^ for C_19_H_33_N_2_O_5_^+^ 369.48, found [M + H]^+^ 369.50.

Tert-butyl ((S)-1-((1R,2S,5S)-2-(((S)-1-amino-1-oxo-3-((S)-2-oxopyrrolidin-3-yl)propan-2-yl)carbamoyl)-6,6-dimethyl-3-azabicyclo[3.1.0]hexan-3-yl)-3,3-dimethyl-1-oxobutan-2-yl)carbamate (**6**). Dipeptide acid **4** (1.22 g, 3.3 mmol) was dissolved in dry CH_2_Cl_2_ (6 mL) and the amine hydrochloride **5** (898 mg, 4.3 mmol), HBTU (1.25 g, 3.6 mmol), and DIPEA (1.4 ml, 8.25 mmol) were added at 0 °C, then DMF (3 mL) was added, and the reaction mixture was kept under stirring at RT for 3 hours. The reaction mixture was washed with 1N HCl (x3), saturated aq. NaHCO_3_ (x3), brine (x3), dried over Na_2_SO_4_, filtered, and concentrated under vacuum. The crude was purified by flash chromatography (CHCl_3_/MeOH 5 to10%) to obtain the tripeptide **6** as a white solid (reaction time: 3 hours, yield: 1.3 g, 75%). ^1^H NMR (400 MHz, DMSO-d_6_) δ 9.41 (d, J = 8.6 Hz, 1H), 8.29 (d, J = 8.9 Hz, 1H), 7.55 (s, 1H), 7.31 (s, 1H), 7.03 (s, 1H), 4.43 (d, J = 8.6 Hz, 1H), 4.35 – 4.25 (m, 2H), 3.91 – 3.84 (m, 1H), 3.67 (d, J = 10.4 Hz, 1H), 3.13 (t, J = 9.0 Hz, 1H), 3.06 – 2.97 (m, 1H), 2.45 – 2.34 (m, 1H), 2.14 (dt, J = 10.5, 7.4 Hz, 1H), 1.97 – 1.86 (m, 1H), 1.70 –1.57 (m, 1H), 1.54 – 1.45 (m, 2H), 1.38 (d, J = 7.7 Hz, 1H), 1.10 (s, 3H), 1.05 (s, 9H), 0.98 (s, 9H), 0.84 (s, 3H). MS (ESI) m/z calcd: [M + H]^+^ for C_26_H_44_N_5_O_6_^+^ 522.67, found [M + H]^+^ 522.70.

(1R,2S,5S)-N-((S)-1-cyano-2-((S)-2-oxopyrrolidin-3-yl)ethyl)-3-((S)-3,3-dimethyl-2-(2,2,2-trifluoroacetamido)butanoyl)-6,6-dimethyl-3 azabicyclo[3.1.0]hexane-2-carboxamide (nirmatrelvir). Tripeptide intermediate **6** (300 mg, 0.57 mmol) was dissolved in CH_2_Cl_2_ (3 mL), the solution was cooled to 0 °C, 4N HCl d in 1,4-dioxane (1.5 mL, 5.7 mmol) and the reaction mixture was stirred at RT for 2 hours. Then, the solvent mixture was evaporated in vacuo and the crude was treated with hexane to obtain the desired compound **7** as HCl salt white solid, which was used without further purification in the following step (reaction time: 2 hours, yield: 261 mg, 100%). Intermediate **7** (230 mg, 0.5 mmol) was suspended in dry CH_2_Cl_2_ (2 mL), under an N_2_ atmosphere, and dry pyridine (0.10 mL, 1.43 mmol) was added. After 30 minutes. the resulting mixture was cooled to 0 °C, TFFA (0.08 mL, 0.57 mmol) was added, and the reaction mixture was kept under stirring at RT for 2 hours. Once observed, by TLC monitoring, the complete conversion of the intermediate **7**, anhydrous pyridine (0.18 mL, 2.28 mmol) was added, and the mixture was cooled at −5°C. After 5 minutes, TFFA (0.16 mL, 1.14 mmol) was added and the reaction was kept under stirring at RT for 18 hours. The solvent was removed under vacuum, the resulting crude was diluted with EtOAc, and the organic phase was washed with 0.5 N HCl (x3), saturated aq. NaHCO_3_ (x1), dried over anhydrous Na_2_SO_4_, filtered, and concentrated under vacuum. The crude was purified by preparative HPLC (Shimadzu LC-20AP; column: Sunfire, 5µm, C18, 100 Å, 19 x 100 mm, C18 with TMS endcapping; mobile phase gradient: 30-75, 20 minutes [H_2_O 0,1% HCOOH, MeCN 0,1% HCOOH]; time course: 20 minutes; t_R_ = 7.8 minutes) to afford the target Nirmatrelvir (yield: 115 mg, 40%) as a white solid. ^1^H NMR (600 MHz, DMSO-d_6_) δ 9.43 (d, J = 8.4 Hz, 1H), 9.03 (d, J = 8.6 Hz, 1H), 7.68 (s, 1H), 4.97 (ddd, J = 10.9, 8.6, 5.1 Hz, 1H), 4.41 (d, J = 8.4 Hz, 1H), 4.15 (s, 1H), 3.91 (dd, J = 10.4, 5.5 Hz, 1H), 3.69 (d, J = 10.4 Hz, 1H), 3.17 - 3.11 (m, 1H), 3.04 (td, J = 9.4, 7.1 Hz, 1H), 2.40 (tdd, J = 10.4, 8.4, 4.4 Hz, 1H), 2.14 (ddd, J = 13.4, 10.9, 4.4 Hz, 1H), 2.11 - 2.03 (m, 1H), 1.76 - 1.65 (m, 2H), 1.57 (dd, J = 7.6, 5.5 Hz, 1H), 1.32 (d, J = 7.6 Hz, 1H), 1.03 (s, 3H), 0.98 (s, 9H), 0.85 (s, 3H). MS (ESI) m/z calcd: [M + H]^+^ for C_23_H_33_F_3_N_5_O_4_^+^ 500.53, found [M+H]^+^ 500.40.

The images of ^1^H NMR of nirmatrelvir are reported below.

^1^H NMR of Nirmatrelvir in DMSO-*d_6_* at 25°C

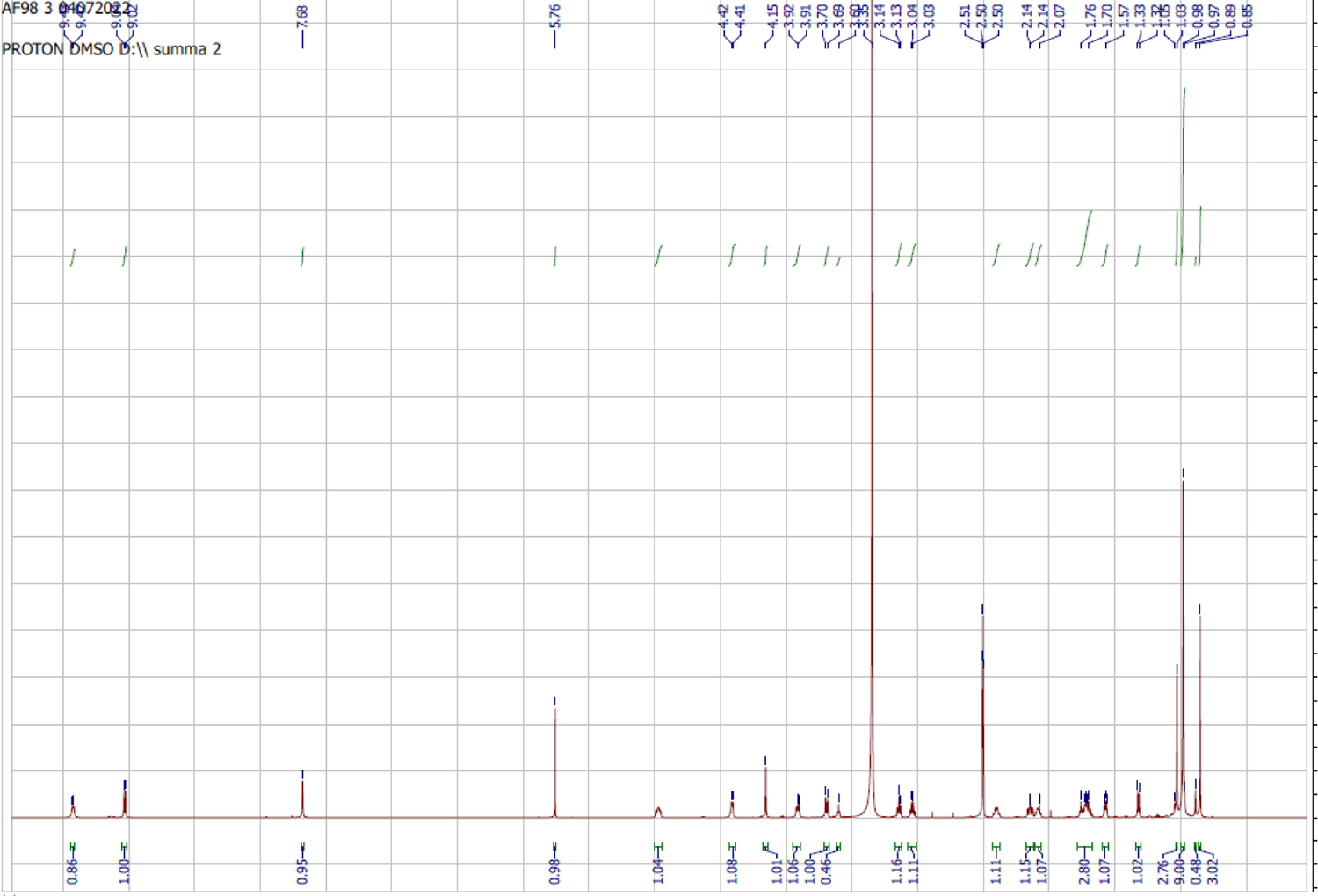

The images of MS spectra of nirmatrelvir are reported below.

Mass spectra of Nirmatrelvir. MS (ESI) *m/z* calcd: [M + H]^+^ for C_23_H_33_F_3_N_5_O_4_^+^ 500.5, found [M+H]^+^ 500.4, [M+Na]^+^ 522.4, [M+K]^+^ 538.3.

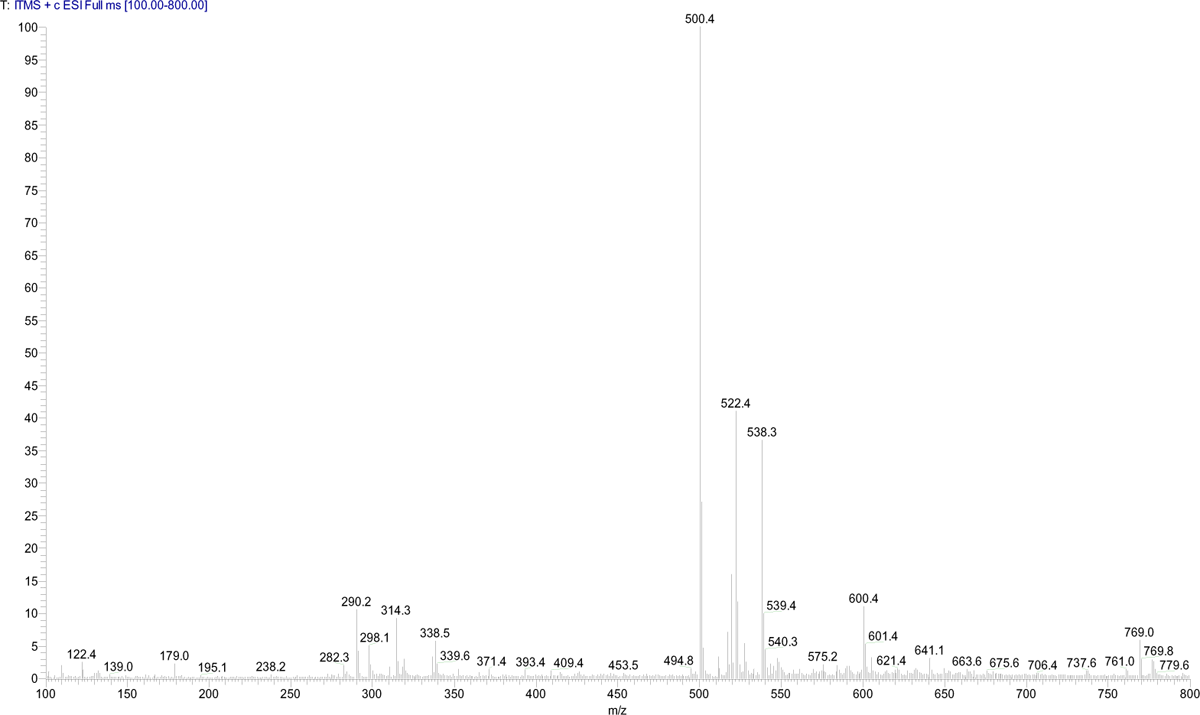

### Biochemical assay

M^pro^ SARS-CoV-2 was expressed in E. coli cells BL21 (DE3) and purified as described (*28, 29*). Briefly, the protein was purified in two steps using a Ni-Sepharose column and by HiTrap Q HP column and the fractions containing the M^pro^ SARS-2 were pooled and concentrated using Amicon Ultra 15 centrifugal filters, at 4000 x g, at 4 °C, in a buffer exchange (20 mM Tris-HCl, 150 mM NaCl, 1 mM EDTA, 1 mM DTT, pH 7.8). Protein purity was verified by SDS-PAGE analysis and the proteins were stored at −80°C.

The M^pro^ SARS-CoV-2 biochemical assays was performed in 386 wells plate in 20 μl of assay buffer containing diluted protein, 20 mM Tris (pH 7.3), 100 mM NaCl and 1mM EDTA, with the addition of 5 mM TCEP, 0.1 % BSA. In Figure 1 **B**, the protein was preincubated for 30 minutes at 37 °C with different concentrations of nirmatrelvir or GC376, a commercially available broad spectrum M^pro^ inhibitor (*17–20*). The substrate DABCYL-KTSAVLQ↓SGFRKM-EDANS (Bachem) was added, and the generation of the fluorescent product was monitored after 15 minutes of incubation (Ex 340 nm, Em 490nm). In **Figure S1A**, the protein and the compound were not preincubated and the enzymatic reaction was immediately initiated with the addition of the substrate in the assay buffer, as described (*4*). The reaction was allowed to progress for 60 minutes at 23° C and then monitored (Ex 340 nm, Em 490nm). Dose response curve were generated by nonlinear regression curve fitting with GraphPad Prism to calculate IC_50_.

### *In vitro* antiviral assays

For *in vitro* antiviral assay HEK293T-hACE were plated in 96 well plates at 5,000 cells/well in complete DMEM plus 2% FBS. After 24 hours, cells were treated with 7 concentration of 5-fold serially diluted nirmatrelvir and infected at 0.1 MOI of SARS-CoV-2 virus. DMSO was used as vehicle for compound serial dilution and not treated control (final concentration 0.25%). Not infected condition was inserted as negative control of infection. Each condition was assayed in three replicates. Antiviral activity was evaluated by qPCR quantification of secreted SARS-CoV-2 RNA and/or by cytopathic effect protection assay (CPE) after 72 hours of incubation at 37°C under 5% CO_2_.

For the quantification of SARS-CoV-2 RNA by qPCR, 10 µl of cell supernatants were subjected to direct lysis with the addition of 10 µl ViRNAex solution (Cabru) and heated at 70°C for 15 minutes. After addition of distilled water (1:1), samples were used as template for PCR amplification using TaqPath™ 1-Step RT-qPCR Master Mix (Thermofisher Fisher Scientific) and specific SARS-CoV-2 primers/probe (2019-nCoV RUO Integrated DNA Technologies). Obtained Ct were normalized to untreated infected wells and dose response curve were generated by nonlinear regression curve fitting with GraphPad Prism to calculate the concentration that inhibits 50% of viral replication (IC_50_).

For CPE assays, CellTiter-Glo® Luminescent Cell Viability Assay (promega), was used. Relative luciferase units (RLUs) were normalized to infected or uninfected controls in order to obtain the percentage of inhibition of cytopathic effect using the following formula: *% CPE inhibition= 100*(Test Cmpd − Avg. Virus)/(Avg. Cells − Avg. Virus),* where *Avg. virus* is the RLU average obtained from infected and not treated wells, while *Avg. Cells* is the RLU average obtained from not infected and not treated wells. Dose response curve were generated by nonlinear regression curve fitting with GraphPad Prism to calculate IC_50_.

### Mice

B6.Cg-Tg(K18-ACE2)^2Prlmn/^J mice (referred to in the text as K18-hACE2) were purchased from The Jackson Laboratory. C57BL/6 mice were purchased from Charles River. Mice were housed under specific pathogen-free conditions and heterozygous mice were used at 8-10 weeks of age. All experimental animal procedures were approved by the Institutional Animal Committee of the San Raffaele Scientific Institute and all infectious work was performed in designed BSL-3 workspaces.

### Mouse infection

Infection of K18-hACE2 transgenic mice with aerosolized SARS-CoV-2 was performed as described (*16*). Briefly, non-anesthetized K18-hACE2 mice were placed in a nose-only Allay restrainer on the inhalation chamber (DSI Buxco respiratory solutions, DSI). To reach a target accumulated inhaled aerosol (also known as delivered dose) of 2 x 10^5^ TCID_50_ mice were exposed to aerosolized SARS-CoV-2 B.1.1.529 or B.1.617.2 for 30-60 minutes (depending on the total volume of diluted virus and on the number of mice simultaneously exposed). In selected experiments, mice were exposed to a target accumulated inhaled aerosol of 1 x 10^6^ TCID_50_. Primary inflows and pressure were controlled and set to 0,5 L/minute/port and −0,5 cmH_2_O, respectively. As control, K18-hACE2 mice received the same volume of aerosolized PBS (125 μL per mouse). Infected mice were monitored daily to record body weight, clinical and respiratory parameters.

C57BL/6 WT mice were infected intravenously with 1 x 10^6^ plaque-forming units (pfu) of VSV Indiana and 2.5 x 10^6^ pfu of LCMV WE.

### *In vivo* treatment

K18-hACE2 mice were treated by oral gavage with nirmatrelvir at 150 mg/kg or vehicle (0,5% Methylcellulose (Methocel A4M, Sigma #94378), 2% Tween80 (Sigma #8170611000) in purified water) for six times starting 4 hours post infection, and every 12 hours thereafter.

### LC-MS/MS Analysis

The nirmatrelvir stock solutions were prepared in DMSO at 1 mg/mL and further diluted to obtain a working solution (WS) at 20 μg/mL. The drug JWH-250 was used as internal standard. The internal standard working solutions (IS-WS) was prepared at 20 ng/mL in methanol:acetonitrile (50:50, v/v) acidified with 0.1 % formic acid.

Plasma of mice were collected, centrifuged at 10,000 rpm for 10 minutes and incubated at 60°C for 30 minutes. The mixture of 30 μL of plasma, 105 μL of IS-WS and 15 μL of WS was vortex for 1 minute and centrifuged at 12,500 g for 10 minutes at 4 °C. The supernatant was collected and 100 μL were injected into the liquid chromatography tandem mass spectrometry (LC–MS/MS) system. The HPLC equipment consisted of an LC AC System from AB Sciex (Toronto, ON, Canada). A Triple Quadrupole Mass Spectrometer (API 2000) from AB-Sciex (Toronto, ON, Canada) was used for detection. The analytes were separated using an Acquity UPLC BEH C18 1.7 μm Column (2.1 x 50 mm ID) from Waters. The mobile phases were (B) MeOH containing 0.2% formic acid and (A) water containing 0.1 % formic acid, at a flow rate of 0.4 mL/min and were entirely transferred into the mass spectrometer source. The gradient elution was as follows: increase of the organic phase from 10 to 100% in 2 minutes and after 1.5 minute of 100 % B the column was led to the original conditions; equilibration of the column was achieved in 2 minutes. Both analytes were detected in positive ionization with a capillary voltage of 4500 V, nebulizer gas (air) at 45 psi, turbo gas (nitrogen) at 70 psi and 450 °C. The other ion source parameters were set as follows: curtain gas (CUR) 25 psi; collision gas (CAD) 6 psi; declustering potential 80 V; entrance potential 8 V. Instrument conditions optimization was performed by direct infusion and manual tuning. Data collection and elaboration were performed by means of Analyst 1.4 software (AB-Sciex). The quantitative data were acquired using Multi Reaction Monitoring (MRM) mode. Two MRM transitions (precursor ion>fragment ion) were selected for the analytes. The parameters used for each analyte are listed in the following table.

|  | Q1 | Q3 | CE (V) | CXP (V) |
| --- | --- | --- | --- | --- |
| Nirmatrelvir | 500.2 | 110.0 | 40 | 16 |
|  |  | 68.0 | 80 | 8 |
| JWH-250 (IS) | 336.1 | 121.0 | 31 | 18 |

The analytical method was validated according to FDA guidelines for bioanalytical method validation. Linearity, precision, accuracy, limits of detection (LODs) and limits of quantification (LOQ) were evaluated. Calibration standard solutions were prepared in blank plasma by spiking 15 μL of a standard mixture at appropriate concentration to 30 μL of plasma and by adding 105 μL of methanol:acetonitrile (50:50, v/v). Calibrators were then treated similarly to the animal samples. The calibration range was 2 to 750 ng/ml and the calibrators were prepared at nine level of concentration. Precision, recovery and accuracy were evaluated at three level of concentrations (25, 100, 750 ng/ml) and resulted within the acceptable limits. LOD was defined as the lowest concentration with a signal-to-noise (S/N) ratio greater than 3. LOQ was defined as the concentration at which both precision (RSD %) and accuracy were less than 20 %. LOQ resulted to be 2 ng/mL while LOD was 1 ng/mL for both analytes.

### Tissue homogenate and viral titers

Tissues homogenates were prepared by homogenizing perfused lungs using gentleMACS Octo Dissociator (Miltenyi BioTec, #130-096-427) in M tubes (#130-093-335) containing 1 ml of DMEM 0% FBS. Samples were homogenized for three times with program m_Lung_01_02 (34 seconds, 164 rpm). The homogenates were centrifuged at 3’500 rpm for 5 minutes at 4°C. The supernatant was collected and stored at −80°C for viral isolation and viral load detection. Viral titer was calculated by 50% tissue culture infectious dose (TCID_50_). Briefly, Vero E6 cells were seeded at a density of 1.5 × 10^4^ cells per well in flat-bottom 96-well tissue culture plates. The following day, 10-fold dilutions of the homogenized tissue were applied to confluent cells and incubated 1 hour at 37°C. Then, cells were washed with phosphate-buffered saline (PBS) and incubated for 72 hours at 37°C in DMEM 2% FBS. Cells were fixed with 4% paraformaldehyde for 30 minutes and stained with 0.05% (wt/vol) crystal violet in 20% ethanol. Limit of detection (LOD) was defined as the lowest concentration whereby the virus, used as positive control, has killing capacity of cells.

### RNA extraction and qPCR

Tissues homogenates were prepared by homogenizing perfused lung and nasal turbinates using gentleMACS dissociator (Miltenyi BioTec, #130-096-427) with program RNA_02 in M tubes (#130-096-335) in 1 ml of Trizol (Invitrogen, #15596018). The homogenates were centrifuged at 2000 g for 1 minutes at 4°C and the supernatant was collected. RNA extraction was performed by combining phenol/guanidine-based lysis with silica membrane-based purification. Briefly, 100 μl of Chloroform were added to 500 μl of homogenized sample and total RNA was extracted using ReliaPrep™ RNA Tissue Miniprep column (Promega, Cat #Z6111). Total RNA was isolated according to the manufacturer’s instructions. qPCR was performed using TaqMan Fast virus 1 Step PCR Master Mix (Lifetechnologies #4444434), standard curve was drawn with 2019_nCOV_N Positive control (IDT#10006625), primer used are: 2019-nCoV_N1-Forward Primer (5’-GAC CCC AAA ATC AGC GAA AT-3’), 2019-nCoV_N1-Reverse Primer (5’-TCT GGT TAC TGC CAG TTG AAT CTG-3’) 2019-nCoV_N1-Probe (5’-FAM-ACC CCG CAT TAC GTT TGG TGG ACC-BHQ1-3’) (Centers for Disease Control and Prevention (CDC) Atlanta, GA 30333). All experiments were performed in duplicate.

### ELISA

Individual sera were titrated in parallel for the presence of SARS-CoV-2 S1 RBD-specific antibody by end-point ELISA. The ELISA plates were functionalized by coating with recombinant Sars-CoV-2 S1 subunit protein (RayBiotech, #230-30162) at a concentration of 2 μg/ml and incubated overnight (O/N) at 4°C. Subsequently, the plates were blocked with 3% fat-free milk, 0,05% Tween20 in PBS for 1 hour at RT. The sera were then added at a dilution of 1/20 (sera from day 7) or 1/500 (sera from day 14, 21, 28) and diluted 1:10 up to 1/1280 or 1/32000, respectively, in duplicate, and the plates were incubated for 2 hour at RT. After five washes with 0,05% Tween20 in PBS, the secondary anti-murine IgG conjugated with horseradish peroxidase (HRP, PerkinElmer, #NEF822001EA) (1:2000) was added and the plates were incubated for 1 hour at RT. After washing, the binding of the secondary was detected by adding the substrate 3,3’,5,5’-tetramethylbenzidine (TMB, BD Biosciences). The reaction was blocked with 0,5 M H_2_SO_4_ and the absorbance at 450 nm and reference 630 nm was measured.

Individual sera were titrated in parallel for the presence of VSV-specific IgG by end-point ELISA. Neutralizing dose 50 were measured as described (*26*).

### SARS-CoV-2 pseudovirus neutralization assay

SARS-CoV-2 pseudovirus neutralization assay was performed as previously described (*30*). Briefly, lentiviral vector containing luciferase reporter were pseudotyped with B.1.1.529 SARS-CoV-2 spike protein and used for neutralization assay. HEK293T-hACE2 receptor were plated in 96 well plates and transduced with 0.05 MOI of SARS-CoV-2 pseudovirus that were subjected to 1 hour at 37°C of preincubation with 3-fold serially diluted mice plasma. After 24 hour of incubation, pseudoparticle cell transduction was measured by luciferase assay using Bright-Glo™ Luciferase System (Promega) and dose response curves were generated by nonlinear regression curve fitting to calculate Neutralization dose 50 (ND50).

### Cell Isolation and Flow Cytometry

Mice were euthanized by cervical dislocation. At the time of autopsy, mice were perfused through the right ventricle with PBS. Nasal turbinates were removed from the nose cavity. Lung tissue was digested in RPMI 1640 containing 3.2 mg/ml Collagenase IV (Sigma, #C5138) and 25 U/ml DNAse I (Sigma, #D4263) for 30 minutes at 37°C. Homogenized lungs were passed through 70 μm nylon meshes to obtain a single cell suspension. Cells were resuspended in 36% percoll solution (Sigma #P4937) and centrifuged for 20 minutes at 2000 rpm (light acceleration and low brake). The remaining red blood cells were removed with ACK lysis. Peripheral blood was collected in PBS 0,5 mM EDTA and lysed two times with ACK. Single-cell suspensions of spleens were generated as described (*26*).

For analysis of *ex-vivo* intracellular cytokine production, 1 mg/ml of brefeldin A (Sigma #B7651) was included in the digestion buffer. All flow cytometry stainings of surface-expressed and intracellular molecules were performed as described (*16, 31–33*). Briefly, cells were stimulated for 4 hour at 37°C in the presence of brefeldin A, monensin (life technologies, #00-4505-51) and a pool of overlapping peptides (1 μg/ml per peptide), including MHC class I- and MHC class II-restricted peptides (9-22 aminoacids) covering the S, S1, S+, M and N protein of SARS-CoV-2 (Miltenyi, #130-126-700; #130-127-041; #130-127-311; #130-126-702, #130-126-698) (*21*). In selected experiments (**Figure S5**), cells were stimulated with GP61 and GP33 peptides (2 μg/ml per peptide) (*34*). As positive control for IFN-γ and TNF-α production, cells were stimulated with PMA (Invitrogen, # 356150050) and Ionomycin (Invitrogen, #I24222). Cell viability was assessed by staining with Viobility™ 405/520 fixable dye (Miltenyi, Cat #130-109-814). In Figure 2 **H**, biotinylated-RDB (26 KDa, kindly provided by L. Aurisicchio from Takis Biotech) were mixed with Alexa Fluor (AF)-647 or 488-fluorescent streptavidins (53 kDa) in a molar ratio of 4:1, respectively, to obtain fluorescent RBD-tetramers at 2 μg/ml. RBD-specific B cells were labeled prior the surface staining for 30 minutes at 4°C.

Antibodies (Abs) are indicated in the table below.

| Name | Clone | Source and catalog number |
| --- | --- | --- |
| CD8 | 53-6.7 | BD Biosciences #558106, Biolegend #100759 |
| CD4 | RM4-5 | Biolegend #100548; BD Biosciences #740208 |
| B220 | RA3-6B2 | BD Biosciences #564662 |
| CD19 | 1D3 | BD Biosciences #749027 |
| CD44 | IM7 | BD Biosciences #741227; BioLegend #103028 |
| CD69 | H1.2F3 | BD Biosciences #612793 |
| CD19 | 1D3 | BD Biosciences #749027 |
| CD62L | MEL-4 | Biolegend #104453 |
| CD44 | IM7 | BD Biosciences #741227; BioLegend #103028 |
| CD45 | 30-F11 | Biolegend #103113, BD Biosciences #564279 |
| IFN-γ | XMG1.2 | BD Biosciences #557735 |
| TNF-α | MP6-XT22 | Biolegend #506329 |
| CD279 (PD-1) | RMP1-30 | BD Biosciences #749306 |
| Granzyme-B | GB12 | Invitrogen #MHGB04 |
| CD86 | GL1 | BD Biosciences #564199 |
| CD80 | 16-10A1 | Biolegend #104738 |
| CD95 | Jo2 | BD Biosciences #557653 |
| CD127 | SB/199 | BD Biosciences #560733 |
| KLRG1 | 2F1/KLRG1 | Biolegend #138409 |
| GL7 | GL7 | Biolegend # 144612 |
| Bcl6 | K112-91 | BD Biosciences #562401 |
| CXCR5 | 2G8 | Biolegend #145532 |
| T-bet | 4B10 | Invitrogen #25-5825-80 |
| ICOS | C398.4A | Biolegend #313537 |
| Streptavidin-AF647 |  | Invitrogen # S32357 |
| Streptavidin-AF488 |  | Invitrogen #S32354 |

Flow cytometry analysis was performed on BD FACS Symphony A5 SORP, Cytek Aurora and analyzed with FlowJo software (Treestar).

### Confocal Immunofluorescence Histology

Mice were euthanized and perfused transcardially with PBS. One left lobe of the lung was collected and fixed in 4% paraformaldehyde for 16 hours, then dehydrated in 30% sucrose prior to embedding in OCT freezing media (Killik Bio-Optica #05-9801). 20 μm sections were cut on a CM1520 cryostat (Leica) and adhered to Superfrost Plus slides (Thermo Scientific). Sections were permeabilized and blocked in PBS containing 0.3% Triton X-100 (Sigma-Aldrich) and 0,5% BSA followed by staining in permeabilization buffer of Foxp3 / Transcription Factor Staining Buffer Set (eBioscence, # 00-5523-00). Slides were stained for Ki-67 (eBioscence, Clone SolA15, # 56-5698-82) and B220 (Biolegend, Clone RA3-6B2, #103228) over night at RT. Lung sections were washed twice for 5 min and stained with DAPI (Life technologies, #D1360) for 5 min at RT, then washed again and mounted for imaging with FluorSaveTM Reagent (Merck Millipore, #345789). Images were acquired on an SP5 or SP8 confocal microscope with 20x objective (Leica Microsystem). To minimize fluorophore spectral spillover, the Leica sequential laser excitation and detection modality was used.

### Statistical analyses and software

Detailed information concerning the statistical methods used is provided in the figure legends. Flow data were collected using FlowJo Version 10.5.3 (Treestar). Statistical analyses were performed with GraphPad Prism software version 8 (GraphPad). Immunohistochemical imaging analyses were performed with QuPath (Quantitative Pathology & Bioimage 5 Analysis) software. *n* represents individual mice analyzed per experiment. Experiments were performed independently at least twice to control for experimental variation. Error bars indicate the standard error of the mean (SEM). In selected experiments (Figure 1B**, C** and **Figure S1A, B**), error bars indicate the standard deviation (SD). Dose–response curves for IC_50_ values were determined by nonlinear regression. We used Mann-Whitney U-tests to compare two groups with non-normally distributed continuous variables and Kruskal-Wallis non-parametric test or One-way ANOVA test to compare three or more unpaired groups. Normality of data distribution was tested with a Shapiro-Wilk normality test and normality were chosen only when normality could be confirmed for each dataset. We used two-way ANOVA followed by Sidak’s multiple comparisons tests to analyze experiments with multiple groups and two independent variables. Significance is indicated as follows: *p < 0.05; **p < 0.01; ***p < 0.001. Comparisons are not statistically significant unless indicated.

